## Supplementary material for "A neural network model for the evolution of learning in changing environments": S1_Appendix

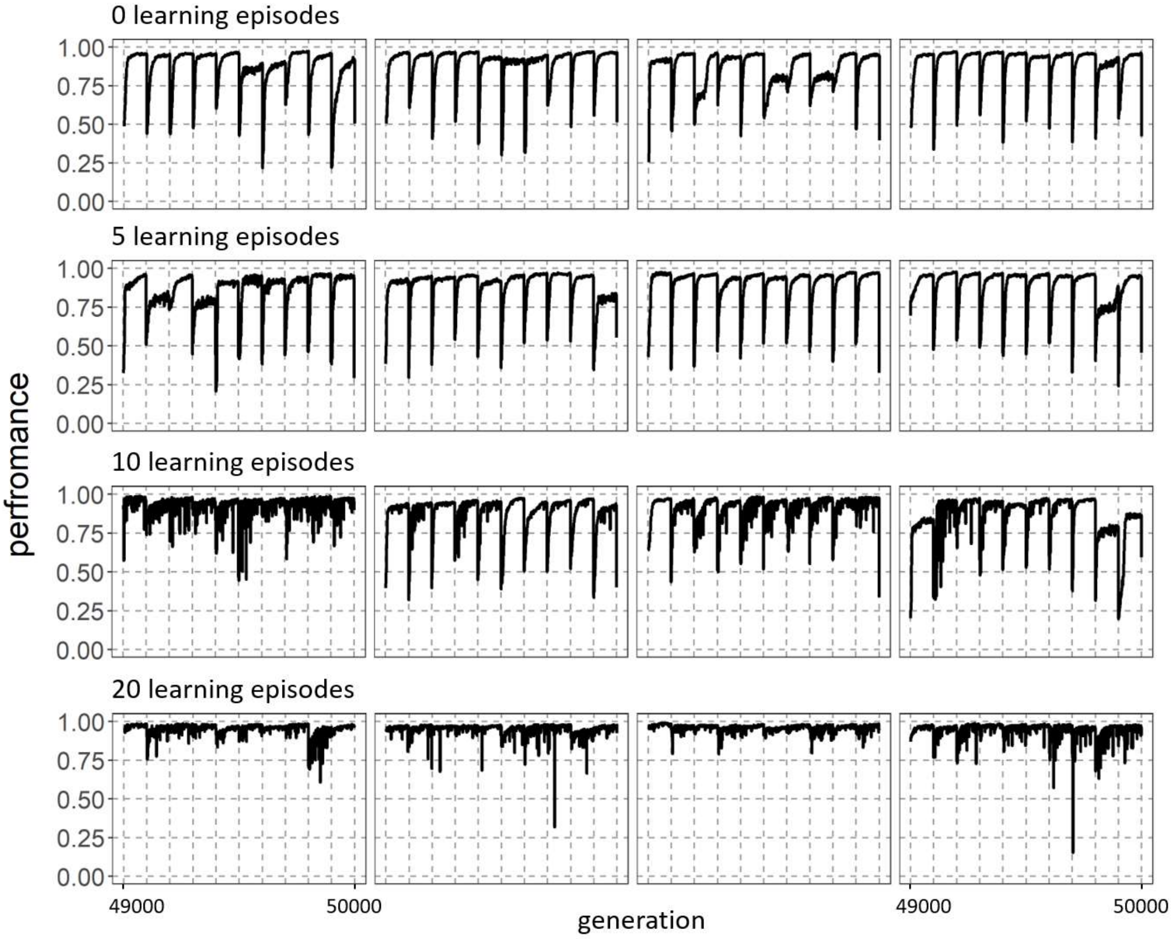


**Fig A. The time course of network performance in a changing environment.** The panels show the time course of average population performance over the last 1000 generations of simulations with the environmental change frequency *f*=0.01 (change once every 100 generations) and magnitude *m*=0.4. For four values of the number of learning episodes (LE = 0, 5, 10, 20) four randomly chosen replicate simulations are shown. It is clear that without learning (LE = 0) or with a short learning period the performance of the population clearly drops every time the environment changes. Only with a large number of learning episodes, performance is better throughout the simulation.

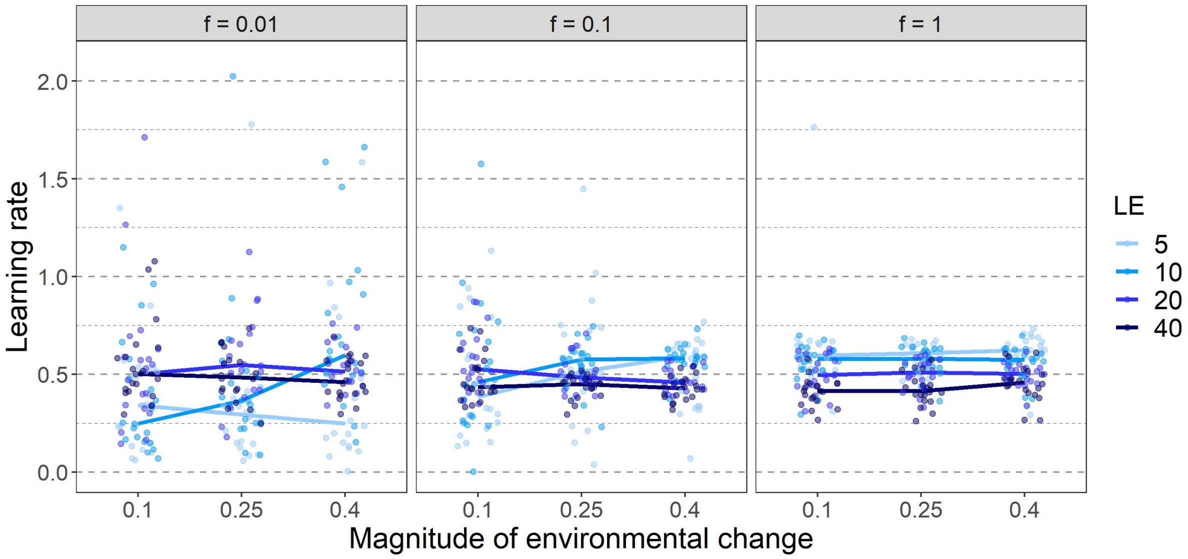


**Fig B. The effect of a fixed number of learning episodes (LE) on the learning rate evolved in different environmental regimes.** Panels in different columns represent different frequencies of environmental change (*f*). The *x*-axis of each panel represents the magnitude of environmental change: the distance that the environmental quality peak moves when change occurs. Points represent the average of the population mean over the last 2000 generations and lines connect medians of all 20 replicates for a given parameter setting. Six data points with values larger than 2.1 are not visible. All of them correspond to LE = 5 (five for frequency of change *f*=0.01, and one for *f*=0.1).

A.

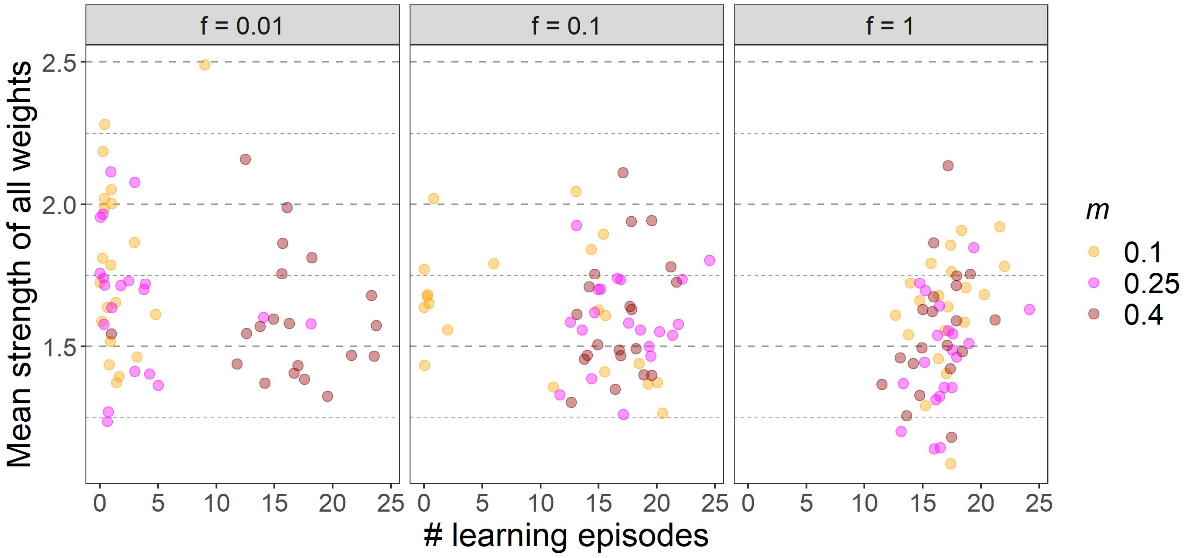


B.

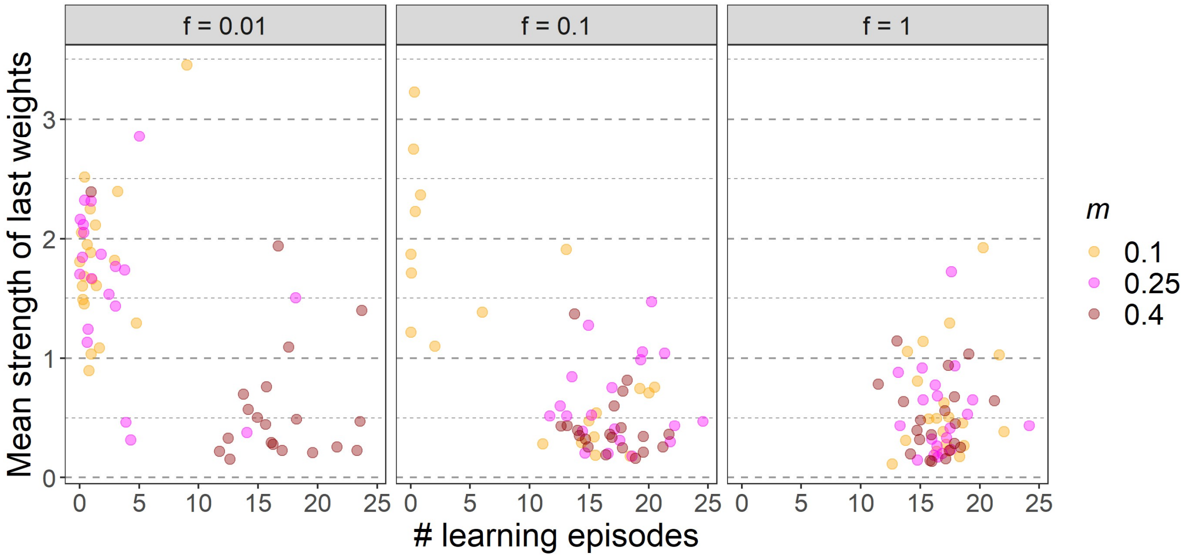


**Fig C. The relationship between weight strength and the number of learning episodes in different environmental regimes. (A)** All weights **(B)** Last, modifiable by learning weights**.** Different columns correspond to different frequencies of environmental change (*f*) and different colours to different magnitudes of environmental change (*m*). Individual mean weight strength was calculated by averaging absolute values of all weights (A) or four weights linked to the output node (B) for each network. Each point represents the population mean over the last 2000 generations of a single replicate. There is no clear correlation between the mean strength of all weights (panel A) with the number of learning episodes. However, the average strength of connections that can be adjusted through learning (panel B) was clearly lower for networks that evolved learning (# learning episodes > 10) compared to networks that did not evolve learning (# learning episodes < 5).

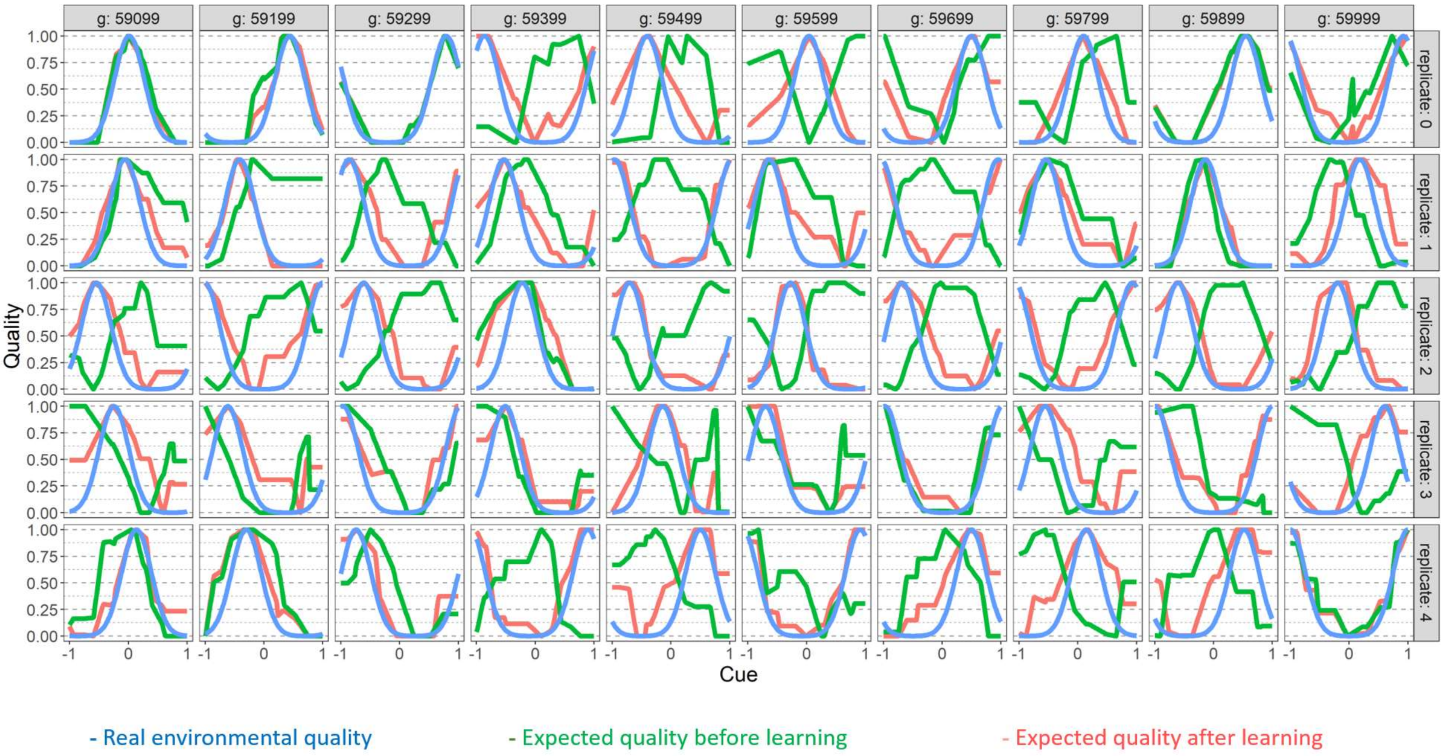


**Fig D. Prediction profiles for evolving networks over time.** The panels show 5 random networks from 5 independent replicates (rows) at different time points (columns) for environmental change of *m*=0.4 every 100 generations (*f*=0.01) and environmental distribution σ = 0.25. Note that the inborn preference (green lines) is relatively stable over time while learning (red) allows for a relatively good matching of predictions with the current environment (blue).

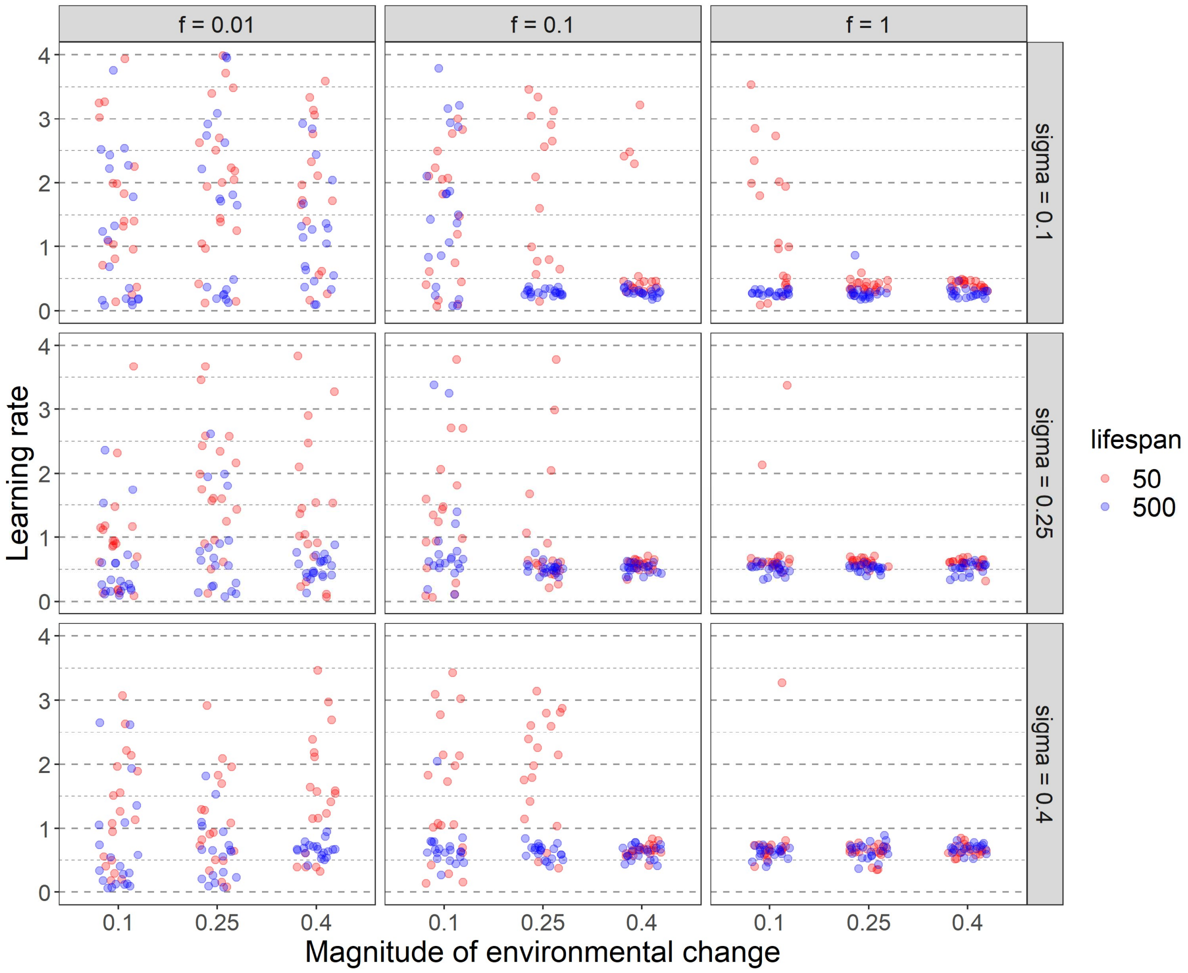


**Fig E. Effect of lifespan and the width of the quality distribution on the evolved learning rate.** This figure corresponds to Fig. 11 in the main text. For the same simulations, it shows how the evolved learning rates depend on the lifespan (50 timesteps: red points, 500 timesteps: blue points) and on the width σ (sigma) of the environmental quality distributions (rows). Graphical conventions as in Fig. 11. The learning rate scale was restricted to 4.0. As a consequence, 37 data points with a learning rate larger than 4.0 are not visible. All of them evolved in replicates without learning, when there is no selection on the learning rate.

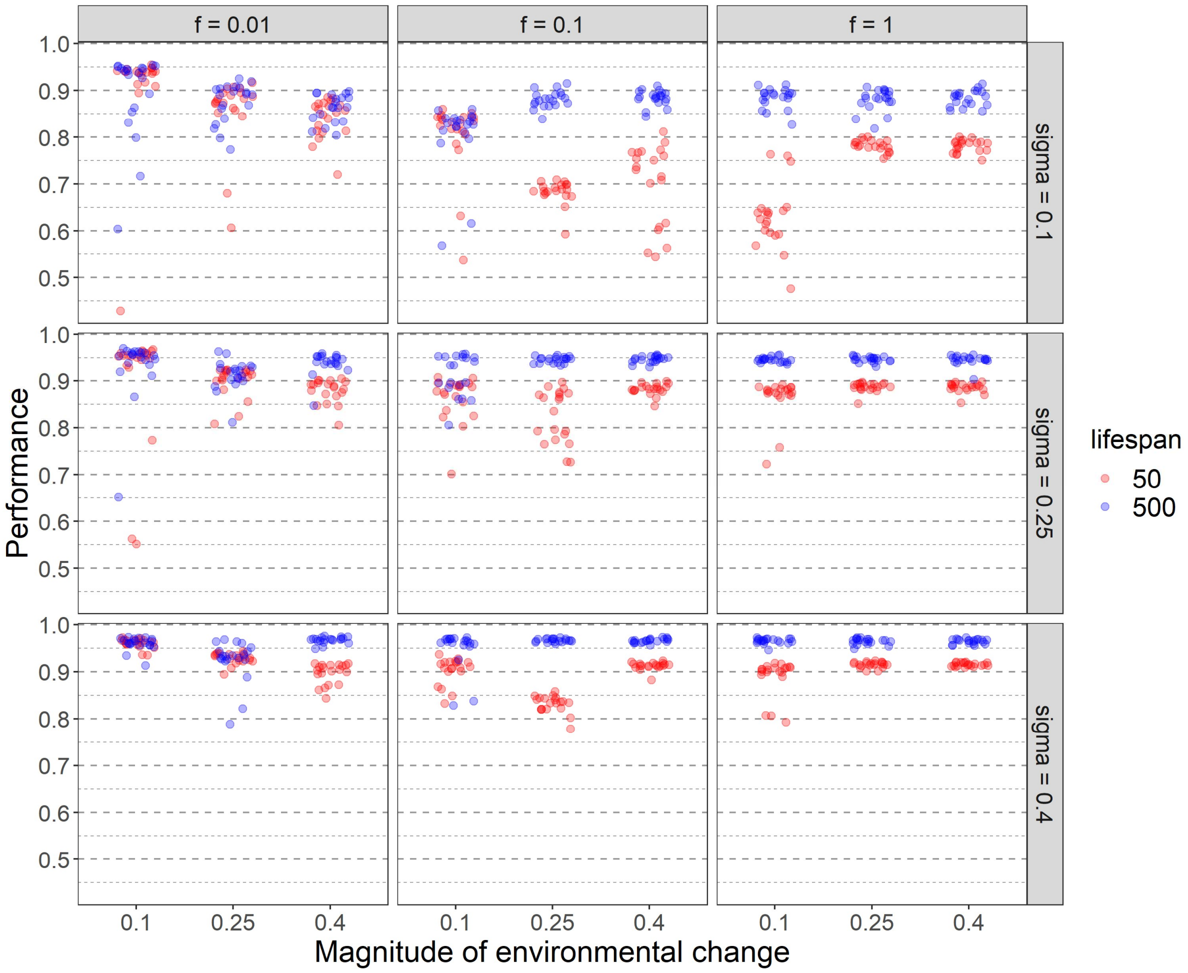


**Fig F. Effect of lifespan and the width of the quality distribution on population performance.** This figure corresponds to Fig E. and Fig. 11 in the main text. For the same simulations, it shows how the average performance in each replicate depends on the lifespan (50 timesteps: red points, 500 timesteps: blue points) and on the width σ (sigma) of the environmental quality distributions (rows). Graphical conventions as in Fig. 11.
